# Prediction of SARS-CoV-2 spike protein mutations using Sequence-to-Sequence and Transformer models

**DOI:** 10.1101/2023.01.23.525130

**Authors:** Hamed Ahmadi, Vahid Nikoofard, Hossein Nikoofard, Rouhollah Abdolvahab, Narges Nikoofard, Mahdi Esmaeilzadeh

## Abstract

In the study of viral epidemics, having information about the structural evolution of the virus can be very helpful in controlling the disease and making vaccines. Various deep learning and natural language processing techniques (NLP) can be used to analyze genetic structure of viruses, namely to predict their mutations. In this paper, by using Sequence-to-Sequence (Seq2Seq) model with Long Short-Term Memory (LSTM) cell and Transformer model with the attention mechanism, we investigate the spike protein mutations of SARS-CoV-2 virus. We make time-series datasets of the spike protein sequences of this virus and generate upcoming spike protein sequences. We also determine the mutations of the generated spike protein sequences, by comparing these sequences with the Wuhan spike protein sequence. We train the models to make predictions in December 2021, February 2022, and October 2022. Furthermore, we find that some of our generated spike protein sequences have been reported in December 2021 and February 2022, which belong to Delta and Omicron variants. The results obtained in the present study could be useful for prediction of future mutations of SARS-CoV-2 and other viruses.

## Introduction

It is well known that SARS-CoV-2 is a virus with single-stranded RNA [1]. RNA viruses have a higher mutation rate than DNA viruses and can adapt themselves to various environmental conditions. The surface of the SARS-CoV-2 is covered by a large number of glycosylated spike proteins that can bind to the receptors on the host cells, Angiotension-Converting-Enzyme2 (ACE2), and mediate viral cell entry [2]. The total length of the SARS-CoV-2 spike protein sequence has 1273 amino acids and comprises a signal peptide (residues 1-13) at the N-terminus, *S*_1_ sub-unit (residues 14-685), and *S*_2_ sub-unit (residues 686-1273) [3, 4]. *S*_1_ sub-unit contains the Receptor-binding-domain which is responsible for the binding of the virus to the cell receptor [5–7]. A virus can have a point mutation, in which another amino acid replaces one amino acid, or amino acids can be deleted. Mutations affect the structural stability and function of the protein. Moreover, it changes the interaction between the spike protein and the ACE2 protein of the cell. For example, mutations of N501, D614, and E484 residues speeds up transmission and infectivity of the SARS-CoV-2 virus [8–10]. Predicting the mutations helps study the evolution of RNA viruses such as influenza, SARS-CoV-2 and related viruses, to develop vaccines with a higher shelf life.

Artificial neural networks are computational models which can be used to study molecular sequence data. Recurrent neural networks (RNNs) are constructive for sequential data. The output in a RNN depends on the prior letters within the sequence and RNN shares parameters across each layer of the network. However, the models based on the RNNs suffer from vanishing and exploding gradient problems when dealing with long data sequences and the precision of the models is reduced [11]. The attention mechanism [12] is an appropriate solution to overcome these problems.

Various works have been done on the applications of neural networks in molecular and computational biology. Neural networks have been used to predict the changes in protein function and its stability due to mutations [**?**, 13–16]. The Tempel model [18] has been used to predict influenza A virus mutation using RNNs, with the attention mechanism. This model can predict whether any mutation happens at a specific site or not. However, the Tempel model provides the probability of mutations and does not provide information about the type of mutations. A previous work [19] constructed a model with 1024 RNN units that could generate spike protein sequences. The dataset used to train this model comprises the spike protein sequences of alpha, beta, gamma and delta coronaviruses. In this model the issue of the temporal evolution of the sequences is not considered.

In this paper, we use the Seq2Seq [20] and the Transformer [21] models to generate the full-length spike protein sequences. The Seq2Seq model maps a sequence to another sequence with the same length or different length and consists of two main RNNs parts. The encoder part reads the input sequence and encapsulates the information as a fixed-length vector called internal state or context vector. The decoder part takes the internal state of the encoder part and uses it for prediction. Without using RNNs, the Transformer model employs the attention mechanism to produce accurate predictions. The attention mechanism improves the performance of the encoder-decoder models in dealing with long data sequences. The attention mechanism finds the parts of the inputs that are important in predicting each instance in the output. The Transformer model can be used in sequential data problems but unlike RNNs, it does not need to process the data in order. Using the self-attention mechanism, parallel computing can speed up training in large datasets.

We generate three sets of the spike protein sequences, 1) by training the models by the dataset from May 2021 to November 2021, 2) by training the models by the dataset from July 2021 to January 2022, and 3) by training the models by the dataset from March 2022 to September 2022. Some of our generated sequences by these models are reported in December 2021 and February 2022. The organization of the paper is as follows. In the next section, preparing of the dataset and the theory of Seq2Seq and Transformer models are described. In Section 3, numerical results of the mutation predictions are presented and discussed. Finally, Section 4 contains a brief summary and conclusion.

## Materials and methods

### Datasets

The global spike glycoprotein of the SARS-CoV-2 sequences which have 1273 amino acids, are collected from the National Center for Biotechnology Information (NCBI) and the duplicate sequences are removed. We cluster sequences according to their months. Visualization of the clusters are shown in supplementary information S1. We use two methods for making datasets. The first one is made by concatenating a random sequence from the first month to another random sequence from the next month. This process continues until the last month. To construct the second dataset, we cluster sequences of each month by using the k-means method [22, 23]. The number of clusters is determined by the elbow method [24]. After clustering the sequences, a sequence from the first month is concatenated to a sequence in the next month in the nearest cluster same as the first dataset. This process continues until the last month. The final result of the two methods is long sequences with the length of *n* × 1273, where *n* is the number of training months. The sequences are made of letters, each one representing an amino acid so they should be transformed into numerical data for feeding the models. For this aim, two methods are used: 1- one-hot encoding [25] for the Seq2Seq model and 2- tokenizing each sequence into a sequence of integers for the Transformer model.

### Sequence-to-Sequence model

The Seq2Seq model is composed of an encoder and a decoder part, which are RNNs of the same type. The encoder-decoder LSTM [26] model may be used to predict the next value in sequential data problems. In the LSTM layer of the encoder by keeping hidden state (*h*) and cell state (*c*), we discard the outputs *y* of the LSTM cells. Here, *y* is a probability distribution over the entire input sequence generated by a softmax activation function. The hidden states for input *x*, are computed as [26]

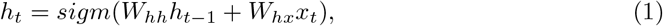

where

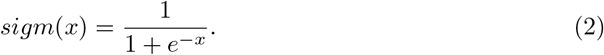

Here *W_hh_*, *W_hx_*, *h_t−1_* and *sigm* are the weight matrix connecting the hidden states, the weight matrix connecting between the hidden states and input, the hidden state of step *t* − 1 and the sigmoid function, respectively [27]. For a sequence with the size of *n*, the LSTM cell reads it in *n* steps and we assign *t*_1_, …, and *t_n_* to specify the steps. The encoder final hidden state (context vector) is calculated from the above equation, which encapsulates the information for all input letters of the sequence. The predicted output at each step is fed as the input in the next step. The output at step *t* is given by the following softmax function [28]

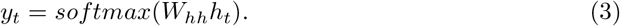

By creating the probability vector, the softmax function determines the final output. We use the teacher-forcing method [29] for training the model. The teacher-forcing method is a quick and efficient training method for RNNs. In this method, the output *y_t_* at step *t* is selected from the training dataset. Then, it is used as the input in the next step in the decoder. The architecture of the Seq2Seq model is shown in Fig S1 in the supplementary information S2.

### Transformer model

Unlike RNN-based models, the Transformer model processes data in parallel to reduce training time for large datasets. The encoder has several identical layers and each layer has two sublayers. The first sublayer works as a multi-head self-attention unit, and the second sublayer is a feed-forward network. The self-attention sublayer reads the input sequence and encodes each letter of the input sequence. Then, the outputs are fed to the feed-forward sublayer network. The output of the final encoder unit is sent to all the decoders (see Fig S2 in the supplementary information S2).

Since there are no RNNs in this model, to remember the order of letters of the input sequence, a relative position is assigned to each letter of the sequence. The assigned positions depend on the order in the sequence. For this purpose, the two following functions are considered [21]

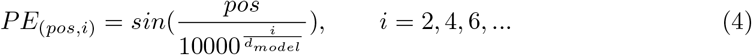

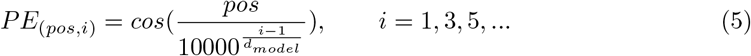

where *pos* and *d_model_* are position and model dimension, respectively. Here, *PE* refers to positional encoding and the model dimension is the dimension of the outputs of the sublayers and the embedding layers of the model. The Transformer model gives attention weights to every token, with the help of the scaled dot-product attention unit. The scaled dot-product attention, which is applied in the model, can be written as [21]

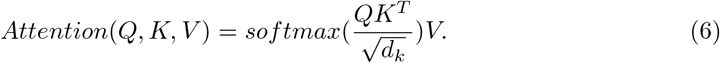

Each attention unit in the model learns the query weights *W_Q_*, the key weights *W_K_*, and the value weights *W_V_*. Each input *x_i_* is multiplied with the weight matrices and produces the query vector *Q_i_* = *x_i_W_Q_*, the key vector *K_i_* = *x_i_W_K_*, and the value vector *V_i_* = *x_i_W_V_*. The attention weight *a_ij_* between tokens *i* and *j* is calculated by the dot product between *Q_i_* and *K_j_*. These attention weights are divided by the square root of the dimension of the key vector, 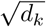. This operation stabilizes gradients in the training process. The attention weights are normalized by using the softmax function. The set of {*W_Q_, W_K_, W_V_*} is an attention head and each layer has multiple attention heads.

The attention function operates in parallel on each set of the projected queries, keys, and values and gives outputs with dimension *d_V_* [21]. Then, the final values are obtained by concatenating and projecting the outputs as follows [21]

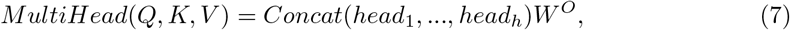

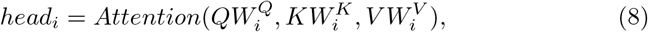

where 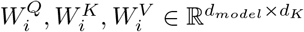, and 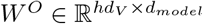.

Feed forward networks in the encoder and the decoder are fully connected networks and operate independently and identically in each position. This operation can be described by the following equation, including two linear transformations and a rectified linear activation function:

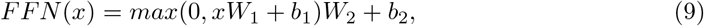

where *b*_1_ and *b*_2_ are the bias vectors and take values during the training process. The linear transformations use different parameters at different layers. The architecture of the Transformer model is shown in Fig S2 in the supplementary information S2.

## Results and Discussion

Using the Transformer and the Seq2Seq models, we generate the spike protein sequences for different months. To generate the spike protein sequences for December 2021, February 2022, and October 2022, we train the models by the random and the clustered datasets, which comprise the spike protein sequences from May 2021 to November 2021, from July 2021 to January 2022, and March 2022 to September 2022, respectively. For Seq2Seq model, we use 5000 training sequences with batch size 8 and 10 epochs, latent dimension 256, Adam optimizer, and accuracy metric. For the Transformer model, we feed the model with 3000 sequences and use the model with a dimension of 128, 8 hidden layers, and 4 heads. Also, we use Adam optimizer and cross-entropy to train the model with batch size 1 and 5 epochs. The models are trained with the randomly connected sequences and the clustered ones. In both scenarios, we reach 99% accuracy. To determine the mutations of the generated spike protein sequences, we compare the generated sequences with the reported SARS-CoV-2 spike protein sequence in Wuhan as a reference sequence. The mutations of the predicted sequences by the models in December 2021, February 2022, and October 2022 are collected in Table 1, Table 2, and Table 3, respectively.

**Table 1.** Spike protein mutations of the predicted sequences by the Seq2Seq and the Transformer models in December 2021.

| Transformer (R)<sup>1</sup> | Seq2Seq (R) | Transformer (C)<sup>2</sup> | Seq2Seq (C) |
| --- | --- | --- | --- |
| L452R | L8del | T19R | L452R |
| T478K | T20del | L452R | T478K |
| D614G | T29del | T478K | D614G |
| P681R | P30del | D614G | P681R |
| D950N | N211del | P681R | D950N |
|  | L452R | D950N |  |
|  | T478K |  |  |
|  | D614G |  |  |
|  | P681R |  |  |
|  | D950N |  |  |

<sup>1</sup>R: Random dataset, <sup>2</sup>C: Clustered dataset

**Table 2.** Spike protein mutations of the predicted sequences by the Seq2Seq and the Transformer models in February 2022.

| Transformer (R)<sup>1</sup> | Seq2Seq (R) | Transformer (C)<sup>2</sup> | Seq2Seq (C) |
| --- | --- | --- | --- |
| A67V | A67V | T19R | T20R |
| H69del | H69del | L452R | R21T |
| V70del | V70del | T478K | T22R |
| T95I | T95I | D614G | L452R |
| N211del | N211del | P681R | T478K |
| L212I | L212I | D950N | D614G |
| ins214 | ins214 |  | P681R |
| G339D | G339D |  | D950N |
| R346K | R346K |  |  |
| S371L | S371L |  |  |
| S373P | S373P |  |  |
| S375F | S375F |  |  |
| D614G | D614G |  |  |
| H655Y | H655Y |  |  |
| N679K | N679K |  |  |
| P681H | P681H |  |  |
| D796Y | D796Y |  |  |
| N856K | N856K |  |  |
| Q954H | Q954H |  |  |
| N969K | N969K |  |  |
| L981F | L981F |  |  |

**Table 3.** Spike protein mutations of the predicted sequences by the Seq2Seq and the Transformer models in October 2022 (Red color: Omicron variant, Blue color: Delta variant).

| Transformer (R) | Seq2Seq (R) | Transformer (C) | Seq2Seq (C) |
| --- | --- | --- | --- |
| F4N | T33G | F2Q | G142D |
| V213G | G35del | D614G | V213G |
| D614G | V36del | D796Y | D614G |
| H655Y | P39D | Q954H | H655Y |
| N679K | D40P | N969K | N679K |
| P681H | V42I |  | P681H |
| Q954H | F43del |  | Q954H |
| N969K | ins50H |  | N969K |
|  | K113S |  |  |
|  | A123T |  |  |
|  | T124del |  |  |
|  | N125del |  |  |
|  | V127I |  |  |
|  | I128V |  |  |
|  | ins133V |  |  |
|  | N137del |  |  |
|  | L141N |  |  |
|  | G142C |  |  |
|  | ins145Y |  |  |
|  | E156del |  |  |
|  | V213G |  |  |
|  | D614G |  |  |
|  | H655Y |  |  |
|  | N679K |  |  |
|  | P681H |  |  |

We search in the NCBI virus database to check whether our generated spike protein sequences have been reported or not. The results are summarized in Table 4 and Table 5. As shown in these tables, among the sequences generated by using the dataset from May 2021 to November 2021, the generated sequence by the Transformer model trained with the random dataset has been reported in USA in December 2021 which belongs to Delta variant (Pango lineage: AY.25). We do not find any reported sequence similar to the generated sequence by the Seq2Seq model trained with the random dataset. The mutations of this generated sequence have been occurred individually, but this set of mutations have not been reported in one spike protein sequence. Also, the generated sequence by the Transformer model trained with the clustered dataset has been reported in December 2021 in USA and belongs to Delta variant (Pango lineage: AY.44). The generated sequence by the Seq2Seq model trained with the clustered dataset belongs to Delta variant (Pango lineage: AY.44), which is reported in December 2021.

**Table 4.** Accession number, Pango lineage, release Date and Location of generated sequences by the Seq2Seq and the Transformer models in December 2021.

|  | Transforme (R) | Seq2Seq (R) | Transformer (C) | Seq2Seq (C) |
| --- | --- | --- | --- | --- |
| Acc<sup>1</sup> | UHT71004.1 | — | UHT71504.1 | UHT76271.1 |
| Pangolin<sup>2</sup> | AY.25 | — | AY.44 | AY.44 |
| Date<sup>3</sup> | 2021-12-29 | — | 2021-12-29 | 2021-12-07 |
| Loc<sup>4</sup> | USA | — | USA | USA |

<sup>1</sup>Acc: Accession number, <sup>2</sup>Pangolin: Pango lineage, <sup>3</sup>Release Date, <sup>4</sup>Location

**Table 5.** Accession number, Pango lineage, release Date and Location of generated sequences by the Seq2Seq and the Transformer models in February 2022.

|  | Transforme (R) | Seq2Seq (R) | Transformer (C) | Seq2Seq (C) |
| --- | --- | --- | --- | --- |
| Acc | ULP59896.1 | ULR99267.1 | UKF77909.1 | — |
| Pangolin | B.1.1.529 | B.1.1.529 | B.1.617.2 | — |
| Date | 2022-02-23 | 2022-02-24 | 2021-12-23 | — |
| Loc | USA | USA | USA | — |

Among the sequences generated by using the dataset from July 2021 to January 2022, the generated sequences by the Transformer and the Seq2Seq models trained with the random dataset belong to Omicron variant (Pango lineage: B.1.1.529), which are reported in February 2022 in USA. The generated sequence by the Transformer model trained with the clustered dataset belongs to Delta variant (Pango lineage: B.1.617.2), which is reported in February 2022 in USA. The generated sequence by the Seq2Seq model trained with the clustered dataset has not been reported.

The generated sequences by the Transformer and Seq2Seq models by using the dataset from March 2022 to September 2022 have not been reported yet. In Table 3, the characteristic mutations of Omicron and Delta variants in the generated sequences in October 2022 are shown by red and blue colors, respectively [30, 31].

## Conclusion

In this study, we used the Transformer and the Seq2Seq models to predict the SARS-CoV-2 virus spike protein mutations. The Transformer and the Seq2Seq models can generate complete mutated sequences of the SARS-CoV-2 spike protein. Most of the predicted spike protein sequences of the SARS-Cov-2 have been reported in several months. The models generate the spike protein sequences of Omicron and Delta variants (Variant Of Concern) of the SARS-CoV-2 virus in the months which we choose as the prediction month: December 2021, February 2022, and October 2022. However, for more accurate predictions, the dimension of the models should be increased and more training dataset should be used.

Having relative knowledge about the changes in virus structure can help researchers to control the disease and to develop vaccines. Mutation intrinsically occurs as a result of a mistake during the genome replication, which is impossible to predict. However, various deterministic conditions such as stability of the resulting mutated protein or the sequence-dependent function of the replicating motor protein are determining in the virus evolution. These deterministic conditions give hope that it may be possible to predict the virus evolutions. Another parameter to consider in the analysis is the geographical distribution of the mutated sequences. A mutated virus often transfers to a nearby person. Thus, considering the geographical distribution may improve the results. The results obtained in this work could be useful for prediction of future mutation of SARS-CoV-2 and other viruses. The presented mehod can also be improved to give more exact results.

## Acknowledgments

All authors were supported by Iran National Science Foundation (INSF) under Grant No. 99023108 (https://insf.org/). The funders had no role in study design, data collection and analysis, decision to publish, or preparation of the manuscript.

## Author Contributions

The authors have declared that no competing interests exist.

## Data Availability

All codes and data files are available from the IUST drive (https://drive.iust.ac.ir/index.ph

